# ViCEKb: Vitiligo-linked Chemical Exposome Knowledgebase

**DOI:** 10.1101/2023.10.10.561820

**Authors:** Nikhil Chivukula, Kundhanathan Ramesh, Ajay Subbaroyan, Ajaya Kumar Sahoo, Gokul Balaji Dhanakoti, Janani Ravichandran, Areejit Samal

## Abstract

Vitiligo is a complex disease wherein the environmental factors, in conjuction with the underlying genetic predispositions, trigger the autoimmune destruction of melanocytes, ultimately leading to depigmented patches on the skin. Apart from being susceptible to other autoimmune disorders, the affected patients may face social stigmatization leading to a decreased quality in life. While genetic factors have been extensively studied, the knowledge on environmental triggers remains sparse and less understood. Therefore, a comprehensive understanding of the environemental triggers will not only explain the complex etiology of the disease, it will also enable the prioritization of potential toxic chemicals in human chemcical exposome. Towards this, we present the first comprehensive knowledgbase of vitiligo triggering chemicals compiled from published literature namely, Vitiligo-linked Chemical Exposome Knowledgbase (ViCEKb). ViCEKb involved an extensive and systematic manual effort in curation of published literature and subsequent compilation of 113 unique chemical triggers of vitiligo. ViCEKb standardises various chemical information, and categorizes the chemicals based on their evidences and sources of exposure. Importantly, ViCEKb contains a wide range of metrics necessary for different toxicological evaluations. Moreover, our extensive cheminformatics-based analysis of the ViCEKb chemical space highlighted its diversity and uniqueness in comparison to other skin specific chemical regulatory lists. We also observed that ViCEKb chemicals are present in various consumer products and are not regulated. Additionally, a transcriptomics based analysis of ViCEKb chemical perturbations in skin cell samples highlighted the commonality in their linked biological processes. Overall, we present the first comprehensive effort in compilation and exploration of various chemical triggers of vitiligo. We believe such a resource will enable in deciphering the complex etiology of vitiligo and aid in the characterization of human chemical exposome. ViCEKb is freely available for academic research at: https://cb.imsc.res.in/vicekb.

## 1. Introduction

Vitiligo (also known as leukoderma) is a chronic, often long-lasting autoimmune disease, where the melanocytes are pathologically killed ultimately resulting in depigmented patches on the skin (Rodrigues et al., 2017). The characteristic vitiligo linked skin depigmentation recurs after the cessation of conventional treatments, due to the presence of resident memory T cells present in the lesion (Richmond et al., 2018). Epidemiological studies have shown that people diagnosed with vitiligo are also susceptible to other autoimmune disorders like type 1 diabetes, rheumatoid arthritis, pernicious anemia and Addisson’s disease (Alkhateeb et al., 2003; Laberge et al., 2005). Apart from health effects, its visible and early onset in children and young adults negatively affects their psychosocial development, thereby leading to a decreased quality of life (Lilly et al., 2013; Porter et al., 1986). Therefore, understanding the etiology is important to identify novel treatments for vitiligo.

Vitiligo is a complex disease, wherein environmental factors trigger autoimmune pathways depending on the underlying genetic predispositions (Spritz and Andersen, 2017). The genetic loci and variations associated with vitiligo have been studied extensively through various genome-wide association studies (GWAS) (Jin et al., 2012, 2010; Shen et al., 2016). There have simultaneously been efforts to generate comprehensive gene set databases like Gene and Autoimmune Disease Association Database (GAAD) (Lu et al., 2018) and VitiVar (Gupta et al., 2019) that were compiled from the vast resource of published literature. Meanwhile, the data on environmental triggers of vitiligo remains sparse and is less understood.

The concept of exposome has been proposed to understand and evaluate the life course of environmental exposures of an individual, and how such exposures impact their health (Vineis et al., 2020; Vrijheid, 2014; Wild, 2005). In particular, chemical exposures through various sources, including occupational and daily-use products, have been touted to be potential risk factors for developing vitiligo, and identifying such chemicals can help in further elucidating the mechanism of its pathogenesis (Harris, 2017). Therefore, compiling a comprehensive list of known chemical triggers of vitiligo will immensely aid in this process.

In this direction, we present Vitiligo-linked Chemical Exposome Knowledgebase (ViCEKb), the first comprehensive knowledgebase on potential chemical triggers of vitiligo. We manually curated and systematically compiled information from published literature pertaining to chemical-induced vitiligo. Thereafter, we standardized the chemical names by mapping them to standard databases such as PubChem (https://pubchem.ncbi.nlm.nih.gov) and Chemical Abstracts Service (CAS) registry (https://www.cas.org/cas-data/cas-registry). Additionally, from the published literature, we compiled and standardized the sources of exposure and evidence levels. Further, we have leveraged different regulations and chemical lists, and checked for the presence of chemical triggers of vitiligo. We have organized all the data into an online web resource namely, ViCEKb, and made it freely available for academic research at: https://cb.imsc.res.in/vicekb/.

We employed various computational approaches to analyze the chemical list compiled in ViCEKb. Notably, by leveraging various publicly available resources and chemical regulations, we observed that the ViCEKb chemicals are present in consumer products, but are unregulated. Our cheminformatics-based analysis revealed the diversity and uniqueness of chemicals within ViCEKb. However, by exploiting the skin tissue specific chemical signatures, we observed that such diverse chemicals affect similar cellular processes. In sum, ViCEKb is the first ever resource compiling a comprehensive list of potential chemical triggers of vitiligo identified in the environment. The insights gained from the various computational analyses can aid in the prioritization of toxic chemicals, and moreover, potentially facilitate in the identification of other chemical triggers of vitiligo.

## 2. Methods

### 2.1. Workflow for the compilation of chemical triggers of vitiligo with supporting evidence in published literature

The main aim of this study is to compile potential chemical triggers of vitiligo in the human exposome that have reported evidence in the published literature. To this end, we designed a workflow based on mining published literature to identify reported evidence on chemical exposure leading to vitiligo pathogenesis, and thereafter, compiled a curated list of potential chemicals which can trigger vitiligo in humans (Figure 1).

**Figure 1:**
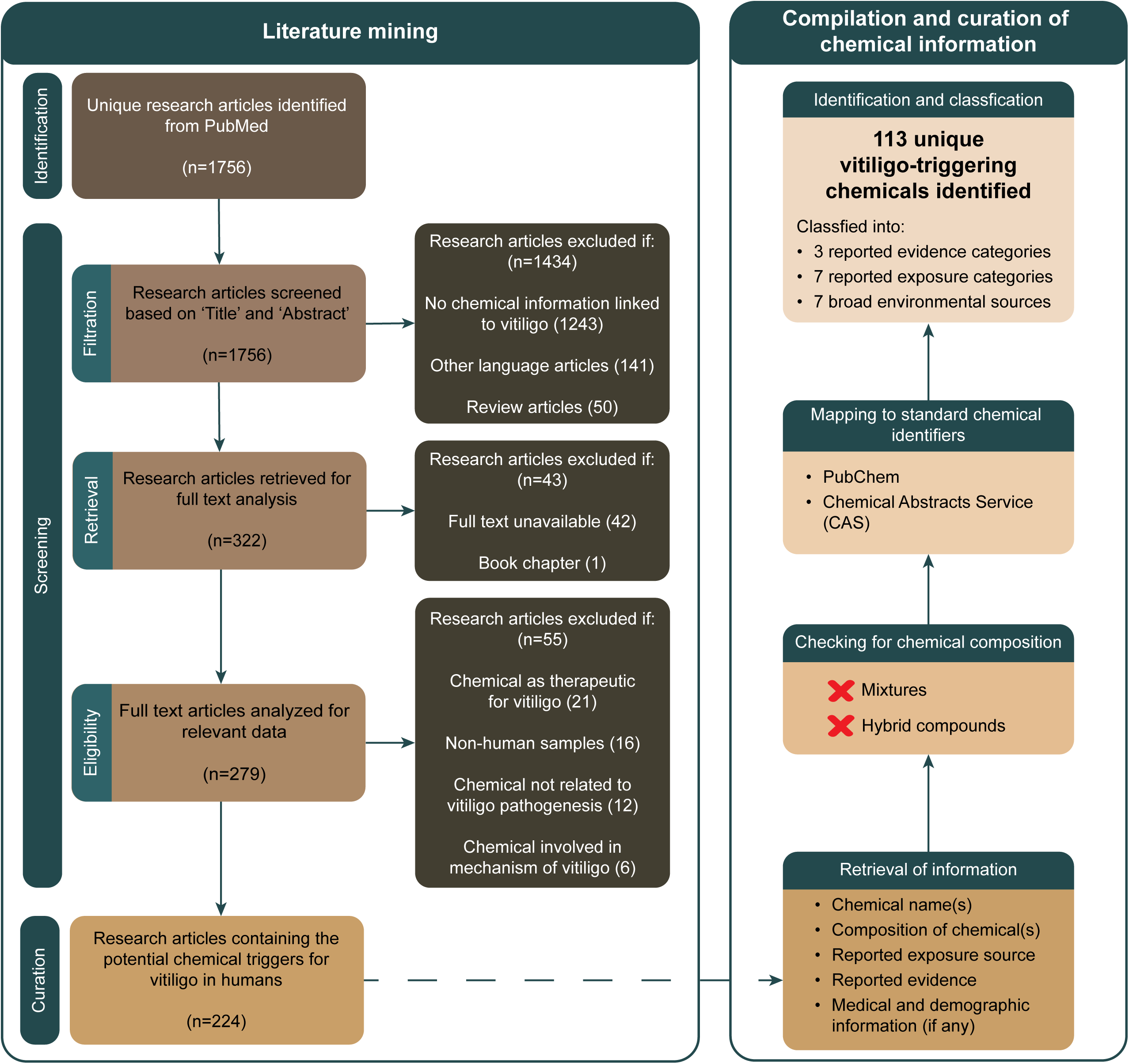
The workflow to identify published literature containing potential chemical triggers of vitiligo is presented as a PRISMA statement. From the shortlisted articles, various chemical information was retrieved and standardized to curate a list of 113 unique chemical-triggers of vitiligo.

First, we mined PubMed (https://pubmed.ncbi.nlm.nih.gov/) using the following query: ‘(leukoderma OR leucoderma OR vitiligo OR hypopigmentation OR depigmentation) AND (chemical* OR exposure* OR expose* OR exposing) AND (human[ORGN] OR “Homo sapiens”[ORGN])’. This PubMed search, last made on 10 August 2022, led to the retrieval of 1756 articles that are likely to contain information on vitiligo-triggering chemicals in humans. Next, we manually screened through the 1756 articles based on their titles and abstracts and excluded articles if: (i) it did not contain any chemical information pertaining to vitiligo pathogenesis; (ii) it is written in a language other than English; (iii) it is a review article (Figure 1). Consequently, we identified 322 articles at the end of the screening process, and thereafter, proceeded to collect their full texts. Here, we excluded an article if it was a book chapter or its full text was unavailable. Accordingly, we retrieved 279 full text articles, following which we manually screened through every article and excluded them if: (i) it contained evidence from non-human organisms; (ii) it reported chemicals involved in the downstream mechanism of vitiligo; (iii) it reported chemical therapeutics for vitiligo; (iv) it reported chemicals not involved in pathogenesis of vitiligo (Figure 1).

Finally, we curated a list of 224 published articles that contained evidence on chemical exposure leading to pathogenesis of vitiligo in humans. The detailed workflow to compile and curate the literature evidence for potential chemical triggers of vitiligo in humans and their corresponding information is presented as a PRISMA statement (Page et al., 2021) in Figure 1. We next describe our workflow to extract the chemical triggers for vitiligo from the 224 published articles.

### 2.2. Curation and compilation of data from shortlisted literature

We manually screened the curated list of 224 published articles to extract information pertaining to potential chemical triggers of vitiligo. From each article, we retrieved the reported chemical name(s), their chemical composition, corresponding chemical exposure source, and the kind of evidence asserting the chemical to be a trigger for vitiligo. We excluded chemical triggers that were reported as multicomponent chemical mixtures because it is difficult to attribute the cause for vitiligo to any of its individual chemical components. Thereafter, we mapped the retrieved chemical names with standard chemical information from PubChem (https://pubchem.ncbi.nlm.nih.gov) and Chemical Abstracts Service (CAS) (https://www.cas.org/cas-data/cas-registry), and subsequently, compiled a list of 113 unique chemicals (Supplementary Table S1). For each of the identified 113 chemicals, we also obtained their identifier from Distributed Structure-Searchable Toxicity (DSSTox) database (Grulke et al., 2019) (https://www.epa.gov/chemical-research/distributed-structure-searchable-toxicity-dsstox-database) to aid in toxicity predictions (Supplementary Table S1). Furthermore, we obtained the chemical categorizations such as chemical Kingdom, Superclass and Class of the identified chemicals using ClassyFire (Djoumbou Feunang et al., 2016) (http://classyfire.wishartlab.com/) (Supplementary Table S1).

### 2.3. Classification of vitiligo triggering chemicals

#### 2.3.1. Based on the type of evidence from published literature

Clinicians and researchers have employed different techniques to associate potential chemical triggers in patients diagnosed with vitiligo (Ghosh and Mukhopadhyay, 2009; Herman et al., 2019; Lee et al., 2016). Based on the supporting evidence used to establish certain chemicals as potential triggers for vitiligo, we broadly classify the identified chemicals into 3 distinct categories. The first category, having the highest level of evidence is ‘Patch test done’, which consists of chemicals that have been tested in small doses on human patient skin and depigmentation was clinically verified. The second category is ‘No patch test done, Observation’ and has a lower level of evidence compared to ‘Patch test done’. This category consists of chemicals inferred from the history of chemical exposure of patients who have been clinically diagnosed with vitiligo. The final category, with the lowest level of evidence is ‘*in vitro* studies’, which consists of evidence from cell lines and enzymatic assays.

#### 2.3.2. Based on the type of chemical exposure

Several sources of chemical exposure have been reported in the published literature (Anthony et al., 2020; Brown et al., 2005; Gawkrodger et al., 1991) which we organize into 7 categories:

i. Cohort study: population-wide studies comprising of various exposure sources
ii. Consumable: chemical exposure through consumption
iii. Cosmetics: chemical exposure through skin care and skin modifying products
iv. Daily-use product: chemical exposure through products used on daily basis
v. Laboratory: *in vitro* chemical studies
vi. Medical: chemical exposure through medical treatments
vii. Occupational: chemical exposure in workplace

Note that chemicals in one category may overlap with those in another category in the above-mentioned categorization.

#### 2.3.3. Based on environmental source

We classified the chemicals based on their reported environmental sources from public documentation curated in PubChem (https://pubchem.ncbi.nlm.nih.gov/). We classified the chemicals into 7 broad categories namely, Agriculture and farming, Consumer products, Industry, Intermediates, Medical and healthcare, Natural Sources and Pollutant, which were further classified into 42 sub-categories (Karthikeyan et al., 2019). Note that a chemical can be classified into multiple categories in the above-mentioned classification.

### 2.4. Physicochemical properties, molecular descriptors and predicted ADMET properties of chemicals

For the identified 113 chemicals, we obtained the respective two-dimensional (2D) and three-dimensional (3D) molecular structures from PubChem. If the 3D structures were not available in PubChem, we employed Open Babel (O’Boyle et al., 2011) to generate the 3D structures from the obtained 2D structures. We employed RDkit (https://www.rdkit.org/) to compute the basic physicochemical properties and the molecular scaffolds of any chemical based on the Bemis-Murcko definition (Bemis and Murcko, 1996). Additionally, we also computed the 1D, 2D and 3D descriptors using PaDEL (Yap, 2011) (http://www.yapcwsoft.com/dd/padeldescriptor/). We have made all this information readily available on our webserver to further aid in the development of chemical toxicity predictors via exploration of the structure-activity relationships.

Absorption, Distribution, Metabolism, Excretion and Toxicity (ADMET) properties can aid in the assessment of chemical toxicity (Shen et al., 2010). We employed several computational tools such as admetSAR 2.0 (Yang et al., 2019), SwissADME (Daina et al., 2017), pkCSM (Pires et al., 2015) and vNN server (Schyman et al., 2017) to predict the ADMET properties for the 113 identified chemicals. The absorption property of chemicals has been estimated based on Caco-2 permeability and skin permeations (log Kp). The potential distribution of the chemical upon absorption has been estimated based on the fraction unbound in human and subcellular localizations. The toxic properties of the chemicals have been estimated based on their cytotoxicity, maximum recommended tolerated dose (MRTD), micronucleus and skin sensitization. For each chemical, we have made these predicted properties readily available on our webserver.

### 2.5. Comparison of vitiligo-triggering chemicals with consumer product database and chemical regulations

Chemicals and Products Database (CPDat) (Dionisio et al., 2018) contains qualitative and quantitative information on the chemical composition of consumer products reported in public domain. Additionally, it provides information on: (i) Organisation for Economic Co-operation and Development (OECD) (https://www.oecd.org/) categorization of functional uses of chemicals; (ii) product compositions and its uses; (iii) regulation in health hazard evaluations. Therefore, we leveraged CPDat to identify the presence of the curated chemical triggers of vitiligo in consumer products.

The CompTox Chemistry Dashboard (Williams et al., 2017) is one of the largest public repositories that aggregates various chemical lists pertaining to toxic effects on humans. Here, we used the skin-specific chemical lists to check if the identified chemicals have been evaluated for their skin sensitizing effects.

Additionally, we checked whether the identified chemicals have been documented for other adverse effects using databases such as DEDuCT (Karthikeyan et al., 2021a) (https://cb.imsc.res.in/deduct/), FCCP (Ravichandran et al., 2022) (https://cb.imsc.res.in/fccp/), ExHuMId (Karthikeyan et al., 2021b) (https://cb.imsc.res.in/exhumid/) and NEUROTOXKB (Ravichandran et al., 2021) (https://cb.imsc.res.in/neurotoxkb/).

### 2.6. Identification of target genes of chemicals triggering vitiligo based on ToxCast skin sensitization assays

The high throughput screening assays undertaken as part of the ToxCast project have helped identify the perturbations in biological processes upon chemical exposure (Richard et al., 2016). Therefore, we leveraged the data within ToxCast version 3.5 (U. S. EPA. 2023, 2022) to identify the potential gene targets of chemicals in human skin samples as follows. First, we checked for the ToxCast assays pertaining to skin tissue and identified 58 relevant assay endpoints (belonging to assay source id 4) from the assay summary file. Then, we filtered the chemical data from level 5 and 6 processing file of assay source id 4 to consider the biologically active (hit_c = 1) representative chemicals (gsid_rep = 1), and identified the necessary chemicals based on their CAS identifiers. Subsequently, we retrieved the target gene, target bioprocess and assay end point information for the identified chemicals.

### 2.7. Exploration of the space of chemicals triggering vitiligo

#### 2.7.1. Chemical similarity network

To assess the similarity of chemicals triggering vitiligo compiled in this study, we constructed a chemical similarity network (CSN) as follows. First, we employed RDKit to obtain the ECFP4 fingerprints of the identified chemicals. Second, we calculated the Tanimoto coefficients (Tanimoto, 1957) between all pairs of chemical fingerprints to check for the similarity between chemicals (Karthikeyan et al., 2019). Third, we constructed a network where nodes are chemicals and edges connect all pairs of nodes with weights given by the computed Tanimoto coefficient between the corresponding pair of chemicals. Next, we filtered out edges that had Tanimoto coefficient < 0.5 and visualized the resulting high similarity backbone of the CSN using Cytoscape (Shannon et al., 2003). Finally, we employed RDKit to compute the maximum common substructure for the connected components identified in the CSN.

#### 2.7.2. Scaffold diversity

In addition to CSN, we probed the similarity between identified chemicals based on their chemical scaffolds. We employed RDKit to compute the scaffolds of the identified vitiligo-triggering chemicals according to the Bemis-Murcko definition (Bemis and Murcko, 1996). To visualize the computed scaffolds, we used the Scopy package (Yang et al., 2021) in python to generate a scaffold cloud representation. Then, we utilized different skin-specific chemical regulatory lists to visualize the scaffold diversity in the identified vitiligo-triggering chemicals. We used RDKit to compute the Bemis-Murcko scaffolds in each of the chemical lists, and analyzed the scaffold diversity via Cyclic System Retrieval (CSR) curves (Vivek-Ananth et al., 2023). Finally, we visualized the scaffold overlaps as an UpSet plot (Lex et al., 2014) using R.

### 2.8. Analysis of transcriptomics signatures of chemicals triggering vitiligo

The Library of Integrated Network-Based Cellular Signatures (LINCS) project has catalogued an extensive dataset of cellular signatures of different human cells upon exposure to various chemical perturbations (Keenan et al., 2018). To analyze the transcriptomics signatures of chemicals triggering vitiligo, we leveraged skin tissue specific L1000 chemical perturbation signatures from iLINCS (Pilarczyk et al., 2022) web-based platform (http://www.ilincs.org/ilincs/). We observed that maximum number of chemicals were tested on cell line A375 for 6 hours at 10μM concentration, and therefore we shortlisted such chemical signatures to ensure maximum chemical coverage. If a chemical has multiple signatures, we selected the latest one containing significantly (p < 0.05) differentially expressed genes. Next, we generated the significantly differentially expressed gene set for each signature by setting the constraints of minimum 2-fold change in expression (|logfoldchange| ≥ 2) and significance value p < 0.05. Thereafter, we employed Enrichr (Chen et al., 2013) (https://maayanlab.cloud/Enrichr/enrich<u>#</u>) to perform gene set enrichment for each of the generated gene sets, and retrieved the enriched pathways from ‘Reactome 2022’ (Gillespie et al., 2022) and enriched biological process ontology terms from ‘GO Biological Process 2023’ (The Gene Ontology Consortium et al., 2023). We additionally set a constraint of adjusted p-value < 0.05, to identify the significantly enriched terms from each of these sets.

### 2.9. Web interface and database management system

We have created an online resource, **Vi**tiligo-linked **C**hemical **E**xposome **K**nowledge**b**ase (ViCEKb), which compiles all the detailed information on the 113 potential chemical triggers of vitiligo identified from 224 published articles. Importantly, ViCEKb provides an easy access to various chemical information such as the type of supporting evidence, environmental sources, chemical structures, physicochemical properties, molecular descriptors, predicted ADMET properties, chemical gene interactions from ToxCast assays, and presence in various skin-specific regulatory lists and chemical exposome databases. ViCEKb is accessible at: https://cb.imsc.res.in/vicekb/.

The data in ViCEKb is stored in MariaDB (https://mariadb.org/) and retrieved using Structure Query Language (SQL). The web interface of ViCEKb was created using PHP (https://www.php.net/) with custom HTML, CSS, jQuery (https://jquery.com/), and Bootstrap 5 (https://getbootstrap.com/docs/5.0/). The interactive charts were created using Google Charts (https://developers.google.com/chart/). ViCEKb is hosted on an Apache (https://www.apache.org) webserver running on Debian 9.4 Linux Operating System.

## 3. Results

### 3.1. ViCEKb – Vitiligo-linked Chemical Exposome KnowledgeBase

Our primary goal is to compile a resource on vitiligo-triggering chemicals within the human exposome. To achieve this, we followed a detailed workflow (Figure 1) to shortlist 224 published articles, from which we compiled and curated the different chemicals names, reported evidence and exposure types (Methods). We identified 113 unique chemicals (Supplementary Table S1) which we categorized under 3 broad evidence categories: ‘Patch test done’ (48 chemicals), ‘No patch test done, Observation’ (53 chemicals) and ‘in vitro studies’ (12 chemicals) (Methods; Figure 2a). We also classified the reported sources of chemical exposure into 7 categories (Methods), of which medical exposure is the most reported source (85 chemicals) (Figure 2b). Additionally, we categorized the chemicals into 7 broad environmental sources (Figure 2c), which we further subdivide into 42 sub-categories (Supplementary Table S2). We observed that the majority of chemicals belong to the environmental source ‘Intermediates’ (80 chemicals) (Figure 2c), which consists of industrial intermediates and human metabolites. We compiled the identified 113 chemicals and associated information in a user friendly, FAIR (**<u>F</u>**indable, **<u>A</u>**ccessible, **<u>I</u>**nteroperable, **<u>R</u>**eproducible) (Wilkinson et al., 2016) and TRUST (**<u>T</u>**ransparency, **<u>R</u>**esponsibility, **<u>U</u>**ser focus, **<u>S</u>**ustainability and **<u>T</u>**echnology) (Lin et al., 2020) compliant webserver named **<u>Vi</u>**tiligo-linked **<u>C</u>**hemical **<u>E</u>**xposome **<u>K</u>**nowledge**<u>b</u>**ase (ViCEKb), which is openly accessible at: https://cb.imsc.res.in/vicekb/.

**Figure 2:**
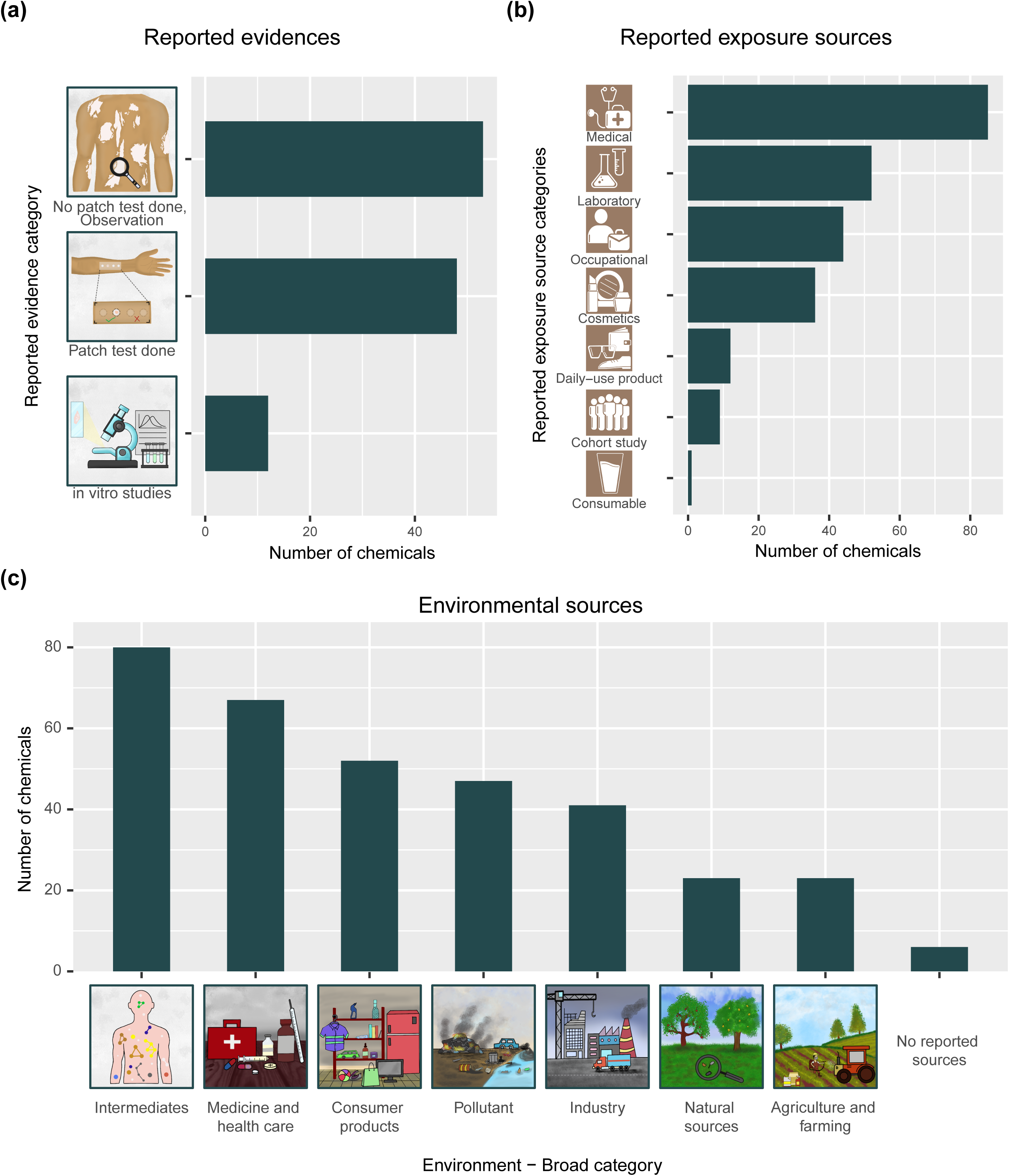
Overview of various chemical information retrieved from published literature. (a) The chemicals have been grouped under 3 categories based on their reported evidence. (b) The chemicals have been grouped under 7 categories based on their reported source of exposure. (c) The chemicals have been grouped under 7 broad environmental categories. Note that 6 chemicals do not have any reported environmental sources, and are therefore unclassified.

The ViCEKb webserver includes various chemical information such as: (i) chemical identifiers; (ii) chemical structures and classification; (iii) chemical properties and descriptors (iv) literature evidence with reported exposure sources; (iv) environmental sources; (v) chemical-gene interactions from ToxCast; (vi) presence of identified chemicals in various regulations including skin-specific regulations and chemical toxicity regulations (Methods). Additionally, it also provides easy access to chemical information compiled from other chemical toxicity and exposome databases like DSSTox (Grulke et al., 2019), DEDuCT 2.0 (Karthikeyan et al., 2021a) (https://cb.imsc.res.in/deduct/), ExHuMId (Karthikeyan et al., 2021b) (https://cb.imsc.res.in/exhumid/), FCCP (Ravichandran et al., 2022) (https://cb.imsc.res.in/fccp/) and NEUROTOXKB (Ravichandran et al., 2021) (https://cb.imsc.res.in/neurotoxkb/).

ViCEKb webserver allows easy navigation through the database with options like BROWSE, SEARCH and DOWNLOAD. The BROWSE option enables users to search for chemicals by their: (i) Evidence level; (ii) Environmental source; (iii) Reported exposure type; (iv) ToxCast skin sensitization target process (Supplementary Figure S1). The SEARCH option allows users to query the chemicals in this database with their names or identifiers - this returns their structure, corresponding evidence from published literature and a link to the chemical information page (Supplementary Figure S2). Additionally, ‘ADVANCED SEARCH’ option provides users with a cheminformatics platform to search chemicals based on physicochemical feature similarity or chemical similarity (Supplementary Figure S2). The datasets in ViCEKb can be downloaded as flat files using the DOWNLOAD option (Supplementary Figure S2). The webserver also has a HELP option that provides a tutorial for using the database.

### 3.2. ViCEKb chemicals are present in consumer products

The pathogenesis of vitiligo has been observed to be linked with the exposure to regular domestic products in developing countries (Ghosh and Mukhopadhyay, 2009). In general, humans depend on a wide variety of consumer products most of which come in direct contact with the skin. Therefore, we explored the presence and functions of chemicals from ViCEKb in consumer products by leveraging CPDat (Dionisio et al., 2018). We identified 91 of 113 chemicals with information on their functional uses, product compositions and their uses, and presence in occupational health hazard reports (Methods), all of which we make available in our webserver under the ‘CPDat information’ section in the chemical information page (Supplementary Figure S3). Among the 91 chemicals, 35 of them were found in various products including personal care and home hygiene (Figure 3a; Supplementary Table S1). Again, 24 of the 91 chemicals had standardized functional uses wherein fragrance was the most reported (Figure 3b; Supplementary Table S1). Additionally, 5 chemicals were found to be documented in health hazard evaluation reports from different industries (Supplementary Table S1). In particular, Isopropyl alcohol (CID:3776) and Hydroquinone (CID:785), which have the highest level of evidence (‘Patch test done’) in ViCEKb, had been documented in health hazard evaluation reports and found in various personal care and hygiene products.

**Figure 3:**
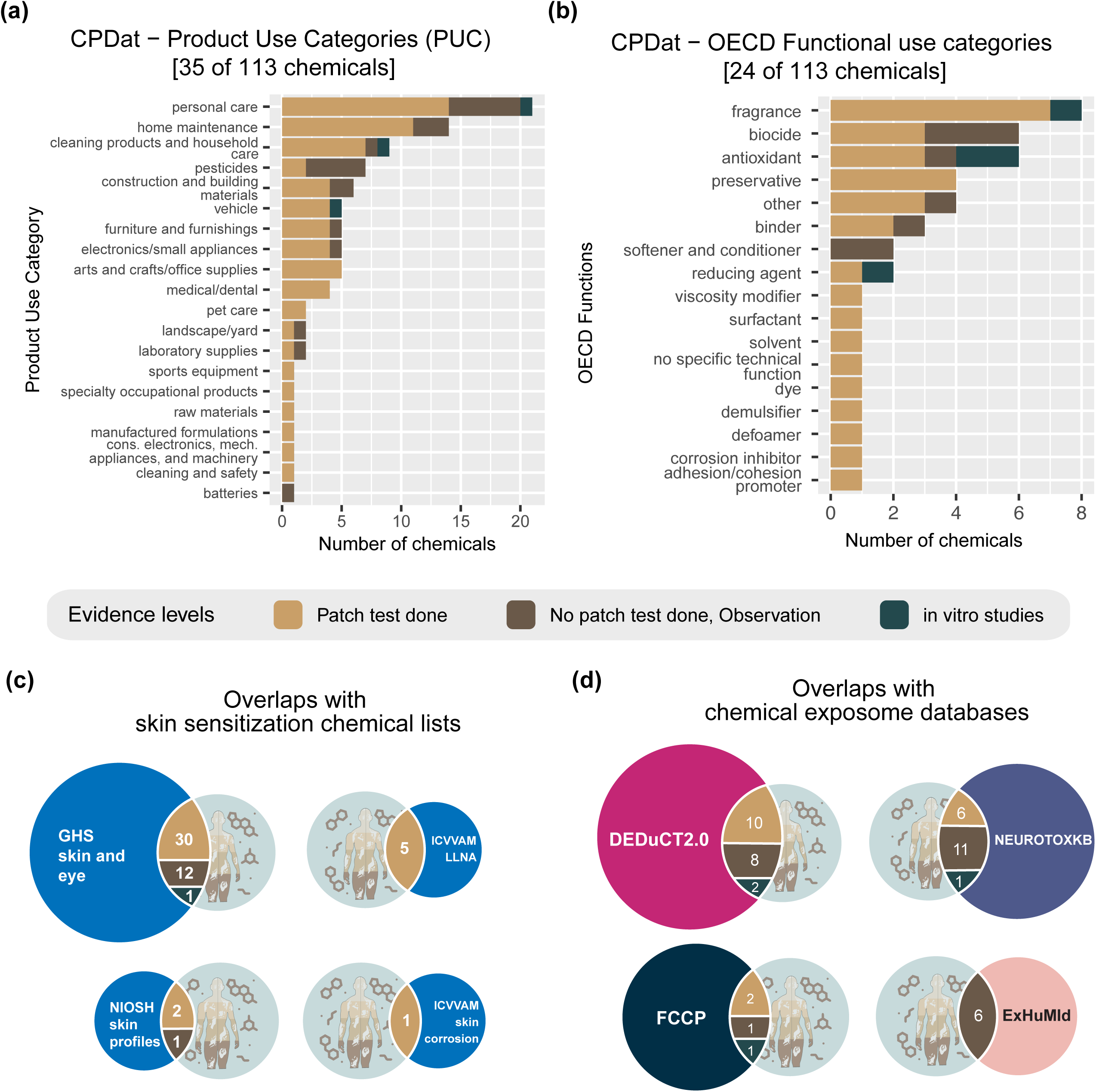
The presence of vitiligo triggering chemicals in consumer products, skin sensitizing chemical lists and other chemical exposome databases. Chemical Products database (CPDat) was leveraged to obtain the consumer product specific data. (a) 35 of 113 chemicals from VICEKb are categorized into 20 product use categories (PUC) in CPDat. (b) 24 of 113 chemicals in ViCEKb are categorized into 17 Organisation for Economic Co-operation and Development (OECD) standardized functions. (c) Number of ViCEKb chemicals overlapping with skin sensitizing chemical lists from CompTox Chemistry Dashboard. (d) Number of ViCEKb chemicals overlapping with chemical exposome databases such as DEDuCT 2.0, NEUROTOXKB, FCCP and ExHuMId. In each of the plots, the chemicals have additionally been classified based on evidence levels in ViCEKb.

### 3.3. ViCEKb shows low overlaps with skin sensitizing chemical lists and exposome databases

The CompTox Chemistry Dashboard (Williams et al., 2017) is one of the largest public domains that aggregates various chemical lists consisting of chemicals evaluated for their toxic effects. The dashboard provides 4 chemical lists pertaining to skin sensitizing chemicals (Supplementary Table S3), and we observed a low overlap with the chemicals in ViCEKb (Figure 3c). Only 2 chemicals with ‘Patch test done’ evidence, namely p-Phenylenediamine (CID:7814) and 2-Mercaptobenzothiazole (CID:697993), have been documented for their skin sensitizing effects in occupational setup. Cinnamaldehyde (CID:637511) was the only chemical with ‘Patch test done’ evidence from ViCEKb prioritized for further laboratory testing for skin corrosion.

Alternatively, to gauge the other adverse effects caused by chemicals in ViCEKb, we leveraged different chemical exposome databases (Methods; Supplementary Table S3), and observed a low overlap (Figure 3d). Among the few overlaps, Hexachlorophene (CID:3598) having ‘Patch test done’ evidence in ViCEKb, is also documented in NEUROTOXKB as a neurotoxicant. Hexachlorophene mainly functions as a preservative and is found in disinfectants and chalkboard cleaners. Another chemical having ‘Patch test done’ evidence in ViCEKb, 4-tert-Butylcatechol (CID:7381) is documented as Category III endocrine disrupting chemical (EDC) in DEDuCT 2.0. 4-tert-Butylcatechol has a variety of functions, including fragrance component in cosmetic products and aiding the production of epoxy resins.

### 3.4. Cheminformatics-based analysis of ViCEKb chemical space revealed its uniqueness

We performed different cheminformatics-based analysis to gain better insights into the chemical space in ViCEKb. First, we characterized the chemicals based on their structures and observed that most chemicals are organic compounds, among which benzenoids are the most prevalant (Methods; Supplementary Table S1). Next, we visualized the similarity among the chemicals in ViCEKb through a chemical similarity network (CSN) and observed that the chemicals are highly diverse with only 28 of the 113 chemicals showing > 50% similarity (Methods; Figure 4; Supplementary Table S4). The high similarity backbone of the CSN shows 12 distinct clusters of chemicals, with the largest clusters containing 6 chemicals each (Methods; Figure 4).

**Figure 4:**
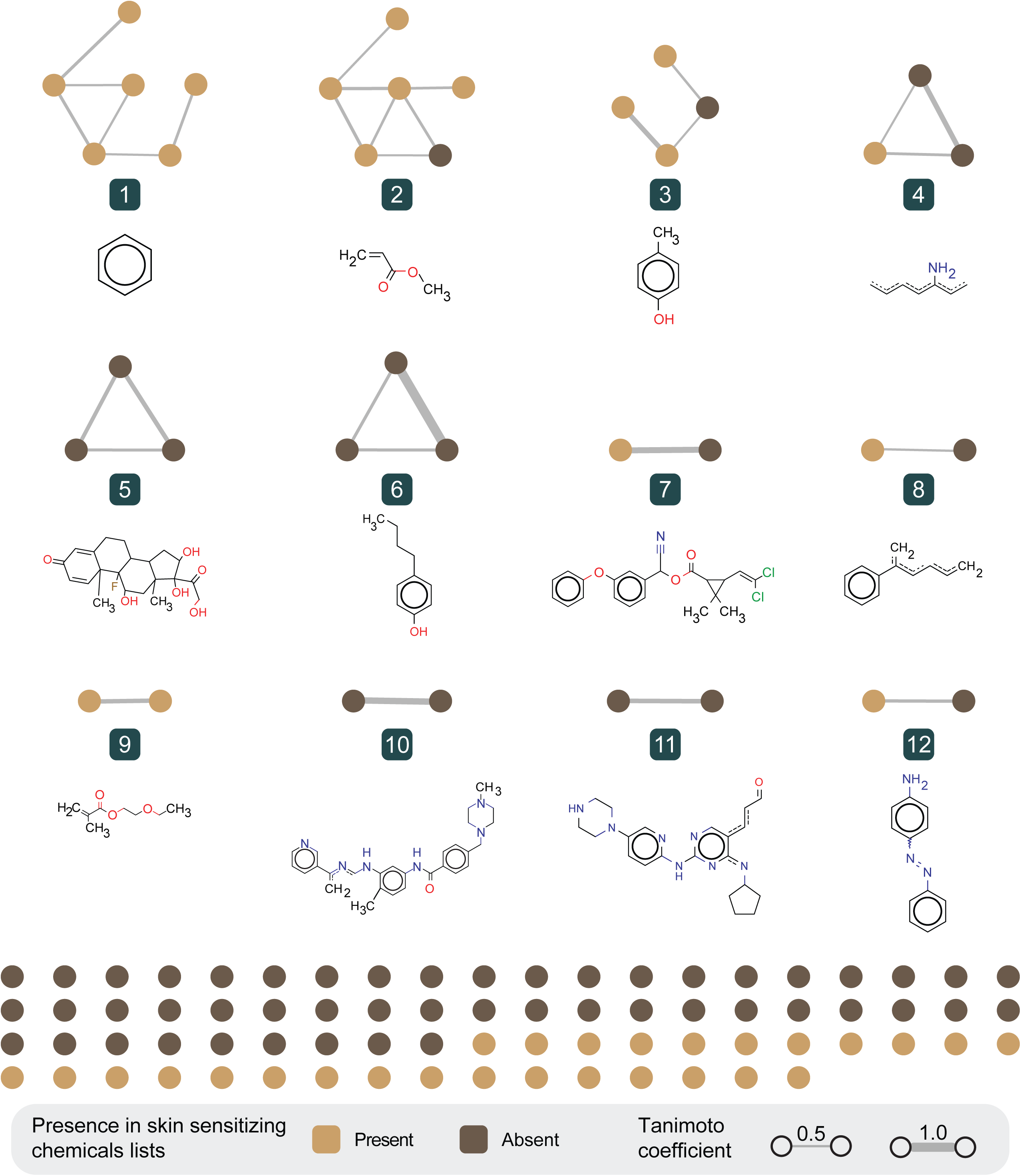
Chemical similarity network of 113 chemicals in ViCEKb. The nodes of the network are chemicals denoted by their presence in the skin sensitizing chemical lists obtained from CompTox Chemistry Dashboard. The edges denote the Tanimoto coefficient computed between the ECFP4 fingerprints of the respective chemicals. The CSN is visualized using Cytoscape 3 software. RDKit was employed to compute the maximum common substructure for each of the 12 connected components identified in the CSN.

Finally, we computed the chemical scaffolds and observed 56 distinct scaffolds across 94 of the 113 chemicals present in ViCEKb (Methods; Figure 5a), where the benzene scaffold was the most represented scaffold (26 of 94) (Supplementary Table S1). Notably, the scaffolds in ViCEKb are the most diverse (AUC – 0.69) compared to those present in cosmetics and skin related regulation lists (Table 1; Figure 5b). Additionally, we observed that 31 of 56 distinct scaffolds in ViCEKb are unique and not captured in these cosmetics and skin related regulations (Figure 5c). These 31 scaffolds span across 32 chemicals, of which 6 chemicals (1,4-Butanediol diglycidyl ether (CID:17046), Neomycin (CID:8378), 2,3,5-Triglycidyl-4-aminophenol (CID:22058861), Thiotepa (CID:5453), Monobenzone (CID:7638), 3-(Diethylamino)-7-(phenylamino)phenoxazin-5-ium (CID:18711204)) have ‘Patch test done’ evidence. Of these chemicals, Neomycin (CID:8378), 2,3,5-Triglycidyl-4-aminophenol (CID:22058861) and 3-(Diethylamino)-7-(phenylamino)phenoxazin-5-ium (CID:18711204) have not been documented in skin sensitizing chemical lists.

**Figure 5:**
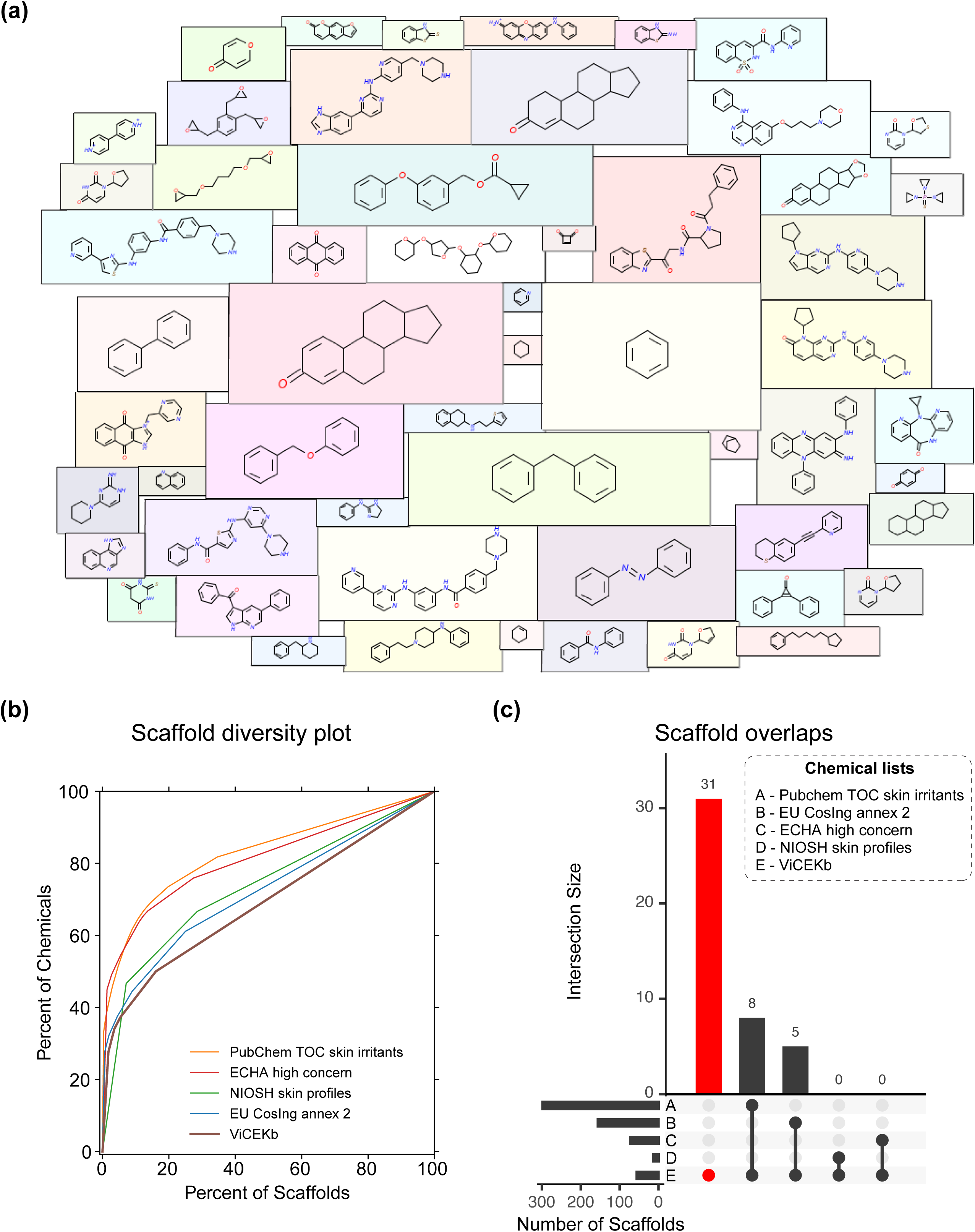
The chemicals in ViCEKb are diverse and unique. (a) A scaffold cloud representation of the 56 unique scaffolds identified from 94 of the 113 chemicals in ViCEKb. The scaffolds are computed using RDKit based on Bemis-Murcko definition. (b) Cyclic System Retrieval (CSR) plot of chemicals in VICEKb, PubChem TOC: skin irritants, ECHA high concern chemicals, NIOSH skin profiles, and EU CosIng annex 2. This plot shows that the ViCEKb is the most diverse in terms of scaffolds. (c) UpSet plot representation of unique overlaps between chemicals in ViCEKb and the other chemical lists, plotted using the UpSetR package in R.

**Table 1:** Comparative analysis of the scaffold diversity of the chemicals in ViCEKb with other chemical lists. Here, M is total number of chemicals in the list, N is the total number of scaffolds in the list, N_sing_ is the total number of singleton scaffolds in the list, AUC is the area under the curve for the corresponding CSR curve, and P_50_ is the percentage of scaffolds required to retrieve 50% of the chemicals in the list.

| | M | N | $N_{\text{sing}}$ | N/M | $N_{\text{sing}}/M$ | $N_{\text{sing}}/N$ | AUC | $P_{50}$ |
| --- | --- | --- | --- | --- | --- | --- | --- | --- |
| ViCEKb | 94 | 56 | 47 | 0.596 | 0.500 | 0.839 | 0.691 | 16.071 |
| NIOSH skin profiles | 30 | 14 | 10 | 0.467 | 0.333 | 0.714 | 0.733 | 14.286 |
| EU CosIng annex 2 | 301 | 156 | 117 | 0.518 | 0.389 | 0.750 | 0.721 | 14.744 |
| ECHA high concern chemicals | 220 | 73 | 53 | 0.332 | 0.241 | 0.726 | 0.810 | 4.110 |
| PubChem TOC: skin irritants | 1069 | 298 | 195 | 0.279 | 0.182 | 0.654 | 0.827 | 4.362 |

**Figure 6:**
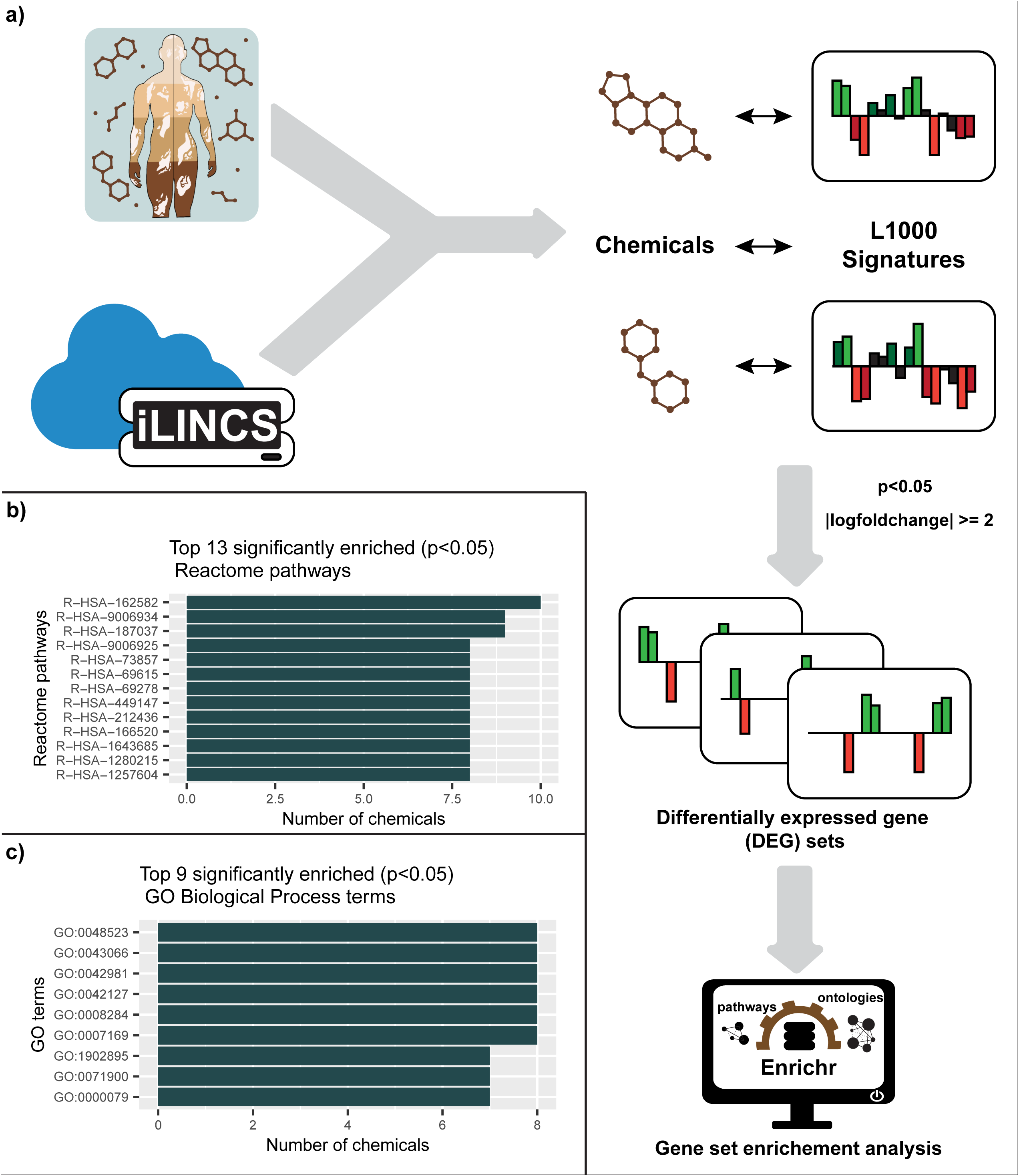
Analysis of transcriptomics signatures of ViCEKb chemicals. (a) The workflow to analyze the significantly enriched pathways and gene ontology (GO) terms from transcriptomics signatures of ViCEKb chemicals. (b) Top 13 significantly enriched reactome pathways present in ViCEKb chemical sets. (c) Top 9 significantly enriched GO biological process terms present in ViCEKb chemical sets.

### 3.5. Vitiligo triggering chemicals are associated with similar biological responses

As the cheminformatics-based analysis of ViCEKb revealed a high structural diveristy among chemcials, we were further interested to explore the nature of their biological responses. A transcriptomics based approach has been used in pharmacology to explore biological similarity between drugs in the context of drug repurposing (Donner et al., 2018; Pilarczyk et al., 2022). By employing a similar approach to identify the similarity among chemicals triggering vitiligo, we obtained skin tissue specific chemical signatures for 18 chemicals, and identified their significantly differentially expressed genes (DEGs) (Methods; Table 2). We performed pathway enrichment analysis for the different sets of DEGs (Methods), and identified that signal transduction pathway (R-HSA-162582) is significantly enriched in 10 of the 18 chemicals (Supplementary Table S5). The gene ontology (GO) enrichement also shows various cellular signaling processes significantly enriched across chemicals (Supplementary Table S6). In particular, receptor tyrosine kinase signaling pathway is significantly enriched across 9 of 18 chemicals. It has been previously observed that chemicals analogous to amino acid tyrosine can interfere in this signaling pathway and result in melanocyte destruction (Harris, 2017), thereby highlighting a biological connection between these 9 chemicals. Notably, these 9 chemicals are categorized under medical exposure, and most of them have been predicted to be highly permeable in skin (Pires et al., 2015).

**Table 2:** L1000 signatures of 18 ViCEKb chemicals obtained from iLINCS web-based platform. The up and downregulated genes are shortlisted using constraints p<0.05 and |logfoldchange| >= 2.

| Chemical | PubChem identifier | LINCS signature identifier | Number of upregulated genes | Number of downregulated genes | Number of DEGs |
| --- | --- | --- | --- | --- | --- |
| Mitoxantrone | CID:4212 | LINCSCP_486 | 124 | 145 | 269 |
| Latanoprost | CID:5311221 | LINCSCP_178719 | 132 | 103 | 235 |
| Sepantronium | CID:10126189 | LINCSCP_2144 | 73 | 110 | 183 |
| Diphenacyprone | CID:65057 | LINCSCP_1913 | 50 | 90 | 140 |
| Palbociclib | CID:5330286 | LINCSCP_2376 | 58 | 61 | 119 |
| Stavudine | CID:18283 | LINCSCP_1948 | 90 | 22 | 112 |
| Dasatinib | CID:3062316 | LINCSCP_2169 | 59 | 50 | 109 |
| Vemurafenib | CID:42611257 | LINCSCP_2385 | 36 | 40 | 76 |
| Triamcinolone Acetonide | CID:6436 | LINCSCP_3096 | 14 | 20 | 34 |
| Triamcinolone | CID:31307 | LINCSCP_2916 | 11 | 11 | 22 |
| Imatinib | CID:5291 | LINCSCP_179309 | 9 | 7 | 16 |
| Nicotin-Amide | CID:936 | LINCSCP_1802 | 7 | 8 | 15 |
| Atomoxetine | CID:54841 | LINCSCP_2866 | 6 | 9 | 15 |
| Methylprednisolone | CID:6741 | LINCSCP_178224 | 5 | 8 | 13 |
| Nevirapine | CID:4463 | LINCSCP_3080 | 4 | 3 | 7 |
| Thiotepa | CID:5453 | LINCSCP_2587 | 5 | 1 | 6 |
| Clofazimine | CID:2794 | LINCSCP_1917 | 0 | 1 | 1 |
| Isotretinoin | CID:5282379 | LINCSCP_2428 | 1 | 0 | 1 |

## 4. Discussion and conclusion

Vitiligo is a complex disease, where both the genetic predispositions and environmental triggers contribute to its pathogenesis. A comprehensive understanding of both the genetic factors and environmental triggers is necessary to understand the complex etiology of vitiligo. While the genetic factors underlying vitiligo have been extensively studied (Gupta et al., 2019; Shen et al., 2016), the environmental factors have been less explored. In this direction, this study presents a first-of-its kind resource on potential chemical triggers of vitiligo with published evidence, namely ViCEKb, which is openly accessible at: https://cb.imsc.res.in/vicekb/.

Building ViCEKB involved an extensive manual effort in systematically curating published literature to identify 113 unique potential chemical triggers of vitiligo, and standardizing their reported evidence and sources of exposure (Figure 1). For each of the 113 potential chemical triggers, ViCEKB compiles the respective chemical information from various standard sources, and contains a wide range of metrics necessary for toxicological evaluations. ViCEKb additionally categorizes the 113 chemicals based on their environmental sources, where maximum chemicals were found to be present as human metabolites under Intermediates (Figure 2c), among which many were also present in medicine and health care products. Notably, medical exposure was also the most reported source of exposure for potential chemical triggers (Figure 2b). Adverse drug reactions (ADRs) refer to the unintended or harmful effects attributed to the use of medicines, and are vital for pharmacovigilance across health sectors (Coleman and Pontefract, 2016; Edwards and Aronson, 2000). The prior knowledge on various side effects of chemicals has aided in the development of different ADR prediction strategies (Xue et al., 2020; Galletti et al., 2022; Park et al., 2023). In this direction, we analyzed the transcriptomics signatures of structurally diverse drugs from ViCEKb, and observed a significant commonality in their linked biological processes. Therefore, we believe ViCEKb with its unique list of chemical triggers of vitiligo can better aid in the prediction of vitiligo triggering drugs.

Further, a cheminformatics-based exploration of the chemical space of ViCEKb revealed its diversity in comparison with different chemical regulation lists (Figure 5). Notably, many of the chemicals are found in a diverse range of consumer products, including cosmetics. Comparison with other chemical exposomes revealed a low overlap of chemicals, revealing the novelty and uniqueness of chemicals in ViCEKb. This highlights the novelty of ViCEKb and provides for identification and prioritization of potential vitiligo triggering chemicals. For instance, propyl gallate (CID:4947) is documented in IFRA transparency list (https://ifrafragrance.org/) as a chemical used in fragrance formulations used in consumer goods. Flutamide (CID:3397) is documented as a safe chemical product in cosmetics (https://cscpsearch.cdph.ca.gov/search/publicsearch), but is found to be an EDC with category III evidence in DEDuCT 2.0. Both these chemicals have the highest level of evidence (‘Patch test done’) in ViCEKb and are predicted to have high permeability through skin (Pires et al., 2015), but are not considered as skin sensitizing chemicals. Additionally, ToxCast skin sensitization assays for both propyl gallate and flutamide reveal downregulation of genes identified as biomarkers (Gupta et al., 2019) in vitiligo pathogenesis.

In conclusion, ViCEKb provides the first comprehensive resource on potential triggers of vitiligo. ViCEKb highlights the presence of unique vitiligo triggering chemicals in daily-use consumer products, which are otherwise overlooked. Moreover, this resource provides extensive chemical information that can enable future efforts in toxicological evaluation of vitiligo triggering chemicals. This resource can additionally aid in the elucidation of the complex etiology of vitiligo, which can in turn aid in proper diagnosis and treatment of this disease.

## Data availability

All the data compiled in this study has been made available in the ‘DOWNLOAD’ section in the associated webserver: https://cb.imsc.res.in/vicekb.

## Acknowledgements

We thank Kishan Kumar for help with figures and illustrations. Areejit Samal would like to dedicate this manuscript to his mother, Suravi Samal, for the inspiration to undertake this study. Areejit Samal acknowledges funding from the Department of Atomic Energy (DAE), Government of India [via Apex project to The Institute of Mathematical Sciences (IMSc), Chennai] and the Max Planck Society, Germany [via a Max Planck Partner Group in Mathematical Biology]. The funders have no role in study design, data collection, data analysis, manuscript preparation or decision to publish.

## CRediT author contribution statement

**Nikhil Chivukula:** Conceptualization, Data Compilation, Data Curation, Formal Analysis, Software, Visualization, Writing; **Kundhanathan Ramesh:** Conceptualization, Data Compilation, Data Curation, Formal Analysis; **Ajay Subbaroyan:** Conceptualization, Data Compilation, Data Curation, Formal Analysis; **Ajaya Kumar Sahoo:** Conceptualization, Data Compilation, Formal Analysis, Software, Visualization; **Gokul Balaji Dhanakoti:** Data Compilation, Data Curation, Software; **Janani Ravichandran:** Conceptualization, Formal Analysis; **Areejit Samal:** Conceptualization, Supervision, Formal Analysis, Writing.

## Supplementary Tables

**Table S1:** The table contains information on the curated list of 113 chemicals. For each chemical, the table provides the chemical identifier, chemical name, Chemical Abstracts Service (CAS) identifier, DSSTox identifier, canonical SMILES, Bemis-Murcko scaffold SMILES, ClassyFire classification (Kingdom, SuperClass, Class), literature identifier(s) (separated by ’|’), ViCEKb evidence level, reported exposure source(s), environmental source sub-category (separated by ’|’), environmental source broad category (separated by ’|’), Chemical and Products Database (CPDat) OECD functional uses (separated by ’|’), CPDat product use compositions (separated by ’|’), CPDat health hazard evaluation reports (separated by ’|’), and gene targets in ToxCast skin sensitization assays (separated by ’|’).

**Table S2:** The table contains information on the 42 environmental sub-categories, along with their broad categories and number of chemicals in this category.

**Table S3:** This table contains the chemical lists used in this work. For each list, the table provides the chemical list name, list description, list type, list source and list accession date.

**Table S4:** This table contains the pairwise chemical similarity values computed as Tanimoto coefficient between ECFC4 fingerprints of the 2 chemicals.

**Table S5:** This table provides the significantly enriched (adjusted p-value<0.05) reactome pathways from the differentially expressed genes across 18 iLINCS signatures. For each pathway, the table provides the list of chemical(s) (separated by ’|’) for which the pathway is significantly enriched.

**Table S6:** This table provides the significantly enriched (adjusted p-value<0.05) GO biological process ontology terms from the differentially expressed genes across 18 iLINCS signatures. For each pathway, the table provides the list of chemical(s) (separated by ’|’) for whcih the term is significantly enriched.

