## Supplementary Figure for "ViCEKb: Vitiligo-linked Chemical Exposome Knowledgebase"

### Supplementary Figures S1-S3

### for

#### ViCEKb: Vitiligo-linked Chemical Exposome Knowledgebase

Nikhil Chivukula[a,b], Kundhanathan Ramesh[a], Ajay Subbaroyan[a,b], Ajaya Kumar Sahoo[a,b], Gokul Balaji Dhanakoti[a], Janani Ravichandran[a,b], Areejit Samal[a,b,*]

[a] *The Institute of Mathematical Sciences (IMSc), Chennai, India*

[b] *Homi Bhabha National Institute (HBNI), Mumbai, India*

**Figure S1:** ViCEKb webserver images. **(a)** The home page of ViCEKb web server with an easy navigation bar. **(b)** Browse for chemicals based on evidence level. **(c)** Browse for chemicals based on environmental categories. **(d)** Browse for chemicals based on reported exposure sources. **(e)** Browse for chemicals based on ToxCast skin sensitization target.

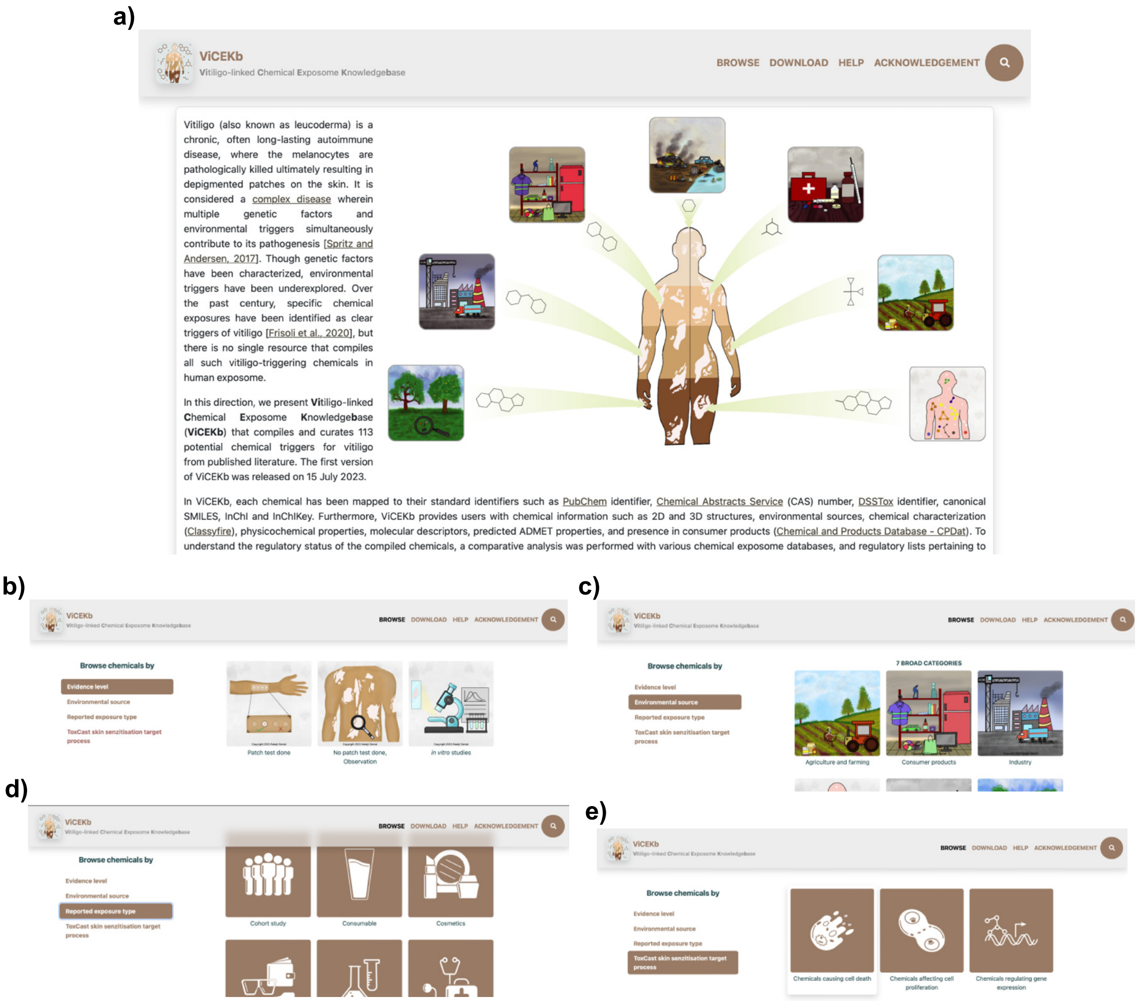

**Figure S2:** ViCEKb webserver images of the 'SEARCH' and 'DOWNLOAD' sections. **(a)** The chemicals can be searched through their chemical identifiers from the top right corner of the page. **(b)** The search results in a tabulated page that contains different information pertaining to the search term. **(c)** ADVANCED SEARCH option based in physicochemical filtrations. **(d)** ADVANCED SEARCH option based on chemical similarity. **(e)** The DOWNLOAD section contains the batch download files compiled in the webserver.

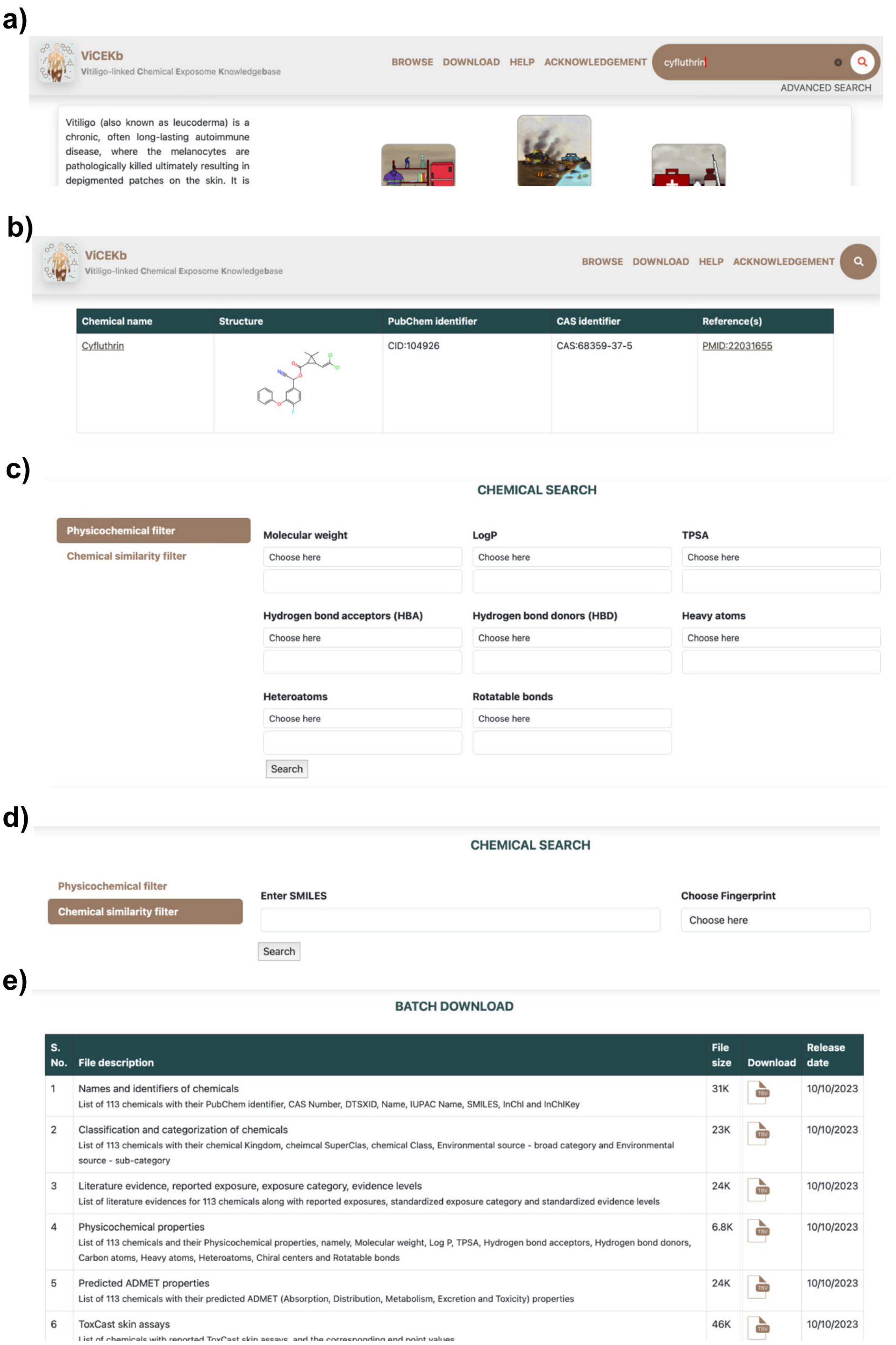

**Figure S3**: ViCEKb webserver images of CHEMCIAL INFORMATION page. **(a)** Chemical identification page containing various chemical information and structure download files. **(b)** CPDat information section containing the various CPDat information divided into different sections.

a)

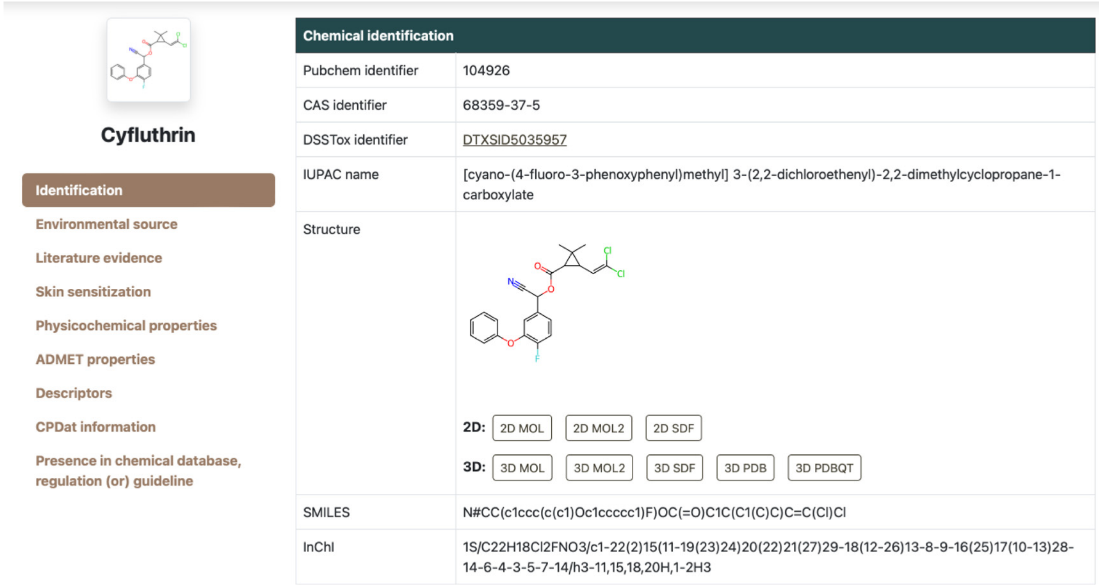
Cyfluthrin

Identification
Environmental source
Literature evidence
Skin sensitization
Physicochemical properties
ADMET properties
Descriptors
CPDat information
Presence in chemical database, regulation (or) guideline

| Chemical identification | |
|---|---|
| Pubchem identifier | 104926 |
| CAS identifier | 68359-37-5 |
| DSSTox identifier | DTXSID5035957 |
| IUPAC name | [cyano-(4-fluoro-3-phenoxyphenyl)methyl] 3-(2,2-dichloroethenyl)-2,2-dimethylcyclopropane-1-carboxylate |
| Structure | 2D: 2D MOL 2D MOL2 2D SDF<br>3D: 3D MOL 3D MOL2 3D SDF 3D PDB 3D PDBQT |
| SMILES | N#CC(c1ccc(c(c1)Oc1ccccc1)F)OC(=O)C1C(C1(C)C)C=C(Cl)Cl |
| InChI | 1S/C22H18Cl2FNO3/c1-22(2)15(11-19(23)24)20(22)21(27)29-18(12-26)13-8-9-16(25)17(10-13)28-14-6-4-3-5-7-14/h3-11,15,18,20H,1-2H3 |

b)

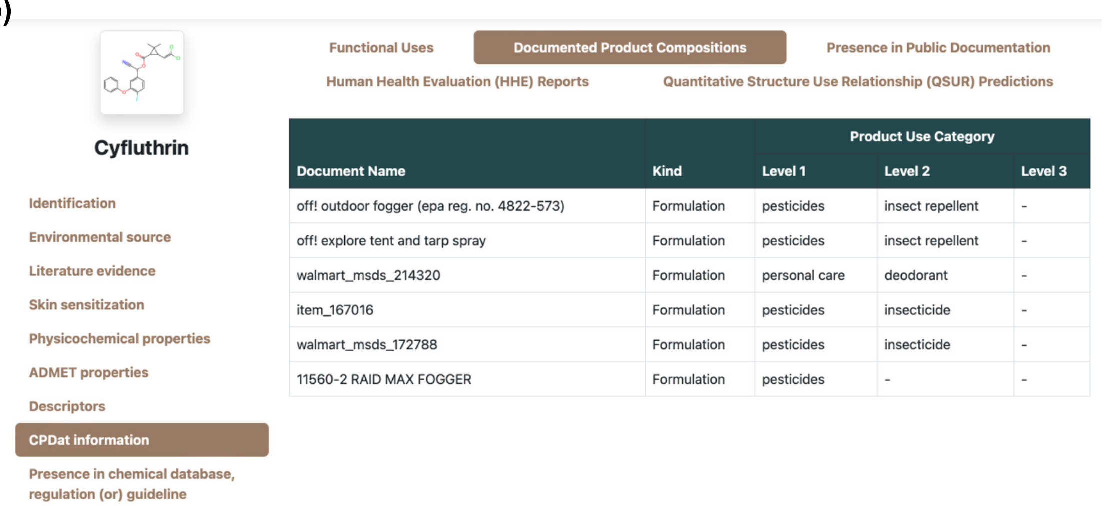
Cyfluthrin

Identification
Environmental source
Literature evidence
Skin sensitization
Physicochemical properties
ADMET properties
Descriptors
CPDat information
Presence in chemical database, regulation (or) guideline

Functional Uses
Documented Product Compositions
Presence in Public Documentation
Human Health Evaluation (HHE) Reports
Quantitative Structure Use Relationship (QSUR) Predictions

| | | Product Use Category | | |
|---|---|---|---|---|
| Document Name | Kind | Level 1 | Level 2 | Level 3 |
| off! outdoor fogger (epa reg. no. 4822-573) | Formulation | pesticides | insect repellent | - |
| off! explore tent and tarp spray | Formulation | pesticides | insect repellent | - |
| walmart_msds_214320 | Formulation | personal care | deodorant | - |
| item_167016 | Formulation | pesticides | insecticide | - |
| walmart_msds_172788 | Formulation | pesticides | insecticide | - |
| 11560-2 RAID MAX FOGGER | Formulation | pesticides | - | - |
